# COSS: A fast and user-friendly tool for spectral library searching

**DOI:** 10.1101/640458

**Authors:** Genet Abay Shiferaw, Elien Vandermarliere, Niels Hulstaert, Ralf Gabriels, Lennart Martens, Pieter-Jan Volders

**Affiliations:** VIB-UGent Center for Medical Biotechnology, VIB, 9000 Ghent, Belgium; Department of Biomolecular Medicine, Ghent University, 9000 Ghent, Belgium; Cancer Research Institute Ghent, Ghent University, 9000 Ghent, Belgium

**Keywords:** tandem mass spectrometry, peptide identification, spectral library searching

## Abstract

Spectral similarity searching to identify peptide-derived MS/MS spectra is a promising technique, and different spectrum similarity search tools have therefore been developed. Each of these tools, however, comes with some limitations, mainly due to low processing speed and issues with handling large databases. Furthermore, the number of spectral data formats supported is typically limited, which also creates a threshold to adoption. We have therefore developed COSS (CompOmics Spectral Searching), a new and user-friendly spectral library search tool supporting two scoring functions. COSS also includes decoy spectra generation for result validation. We have benchmarked COSS on three different spectral libraries and compared the results with established spectral search and sequence database search tool. Our comparison showed that COSS more reliably identifies spectra and is faster than other spectral library searching tools. COSS binaries and source code can be freely downloaded from https://github.com/compomics/COSS.

## INTRODUCTION

Tandem mass spectrometry (MS/MS) is a commonly used method to analyze and identify peptides and proteins. Typically, MS/MS analysis and identification consists of several steps^1^. First, an unknown protein mixture is digested into peptides with the aid of a protease, and the resulting peptides are then separated in time by liquid chromatography (LC). This LC is coupled directly to a mass spectrometer’s source where the eluting peptides are detected, selected, and fragmented. The resulting fragment ions are then analyzed by a second stage of mass spectrometry to acquire an MS/MS spectrum. These MS/MS spectra can then be subjected to different computational approaches to match them to peptide sequences.

Commonly used approaches are *de novo* sequencing, sequence database searching, and spectral library searching. *De novo* sequencing^2^ algorithms directly infer the amino acid sequence from the experimental spectrum. In this technique, the quality of the spectrum affects the success of the inference process and hence the identification result. Therefore, the identification rate of such algorithms in practice is typically limited^3^, in turn limiting their use. In sequence database searching, an *in silico* digest of a protein sequence database produces a list of peptide sequences, each of which is then used to generate theoretical mass spectra. These theoretical spectra are subsequently compared with experimental spectra using a similarity scoring function. Due to their performance, sequence database search engines are the most widely used approach to analyze MS/MS data. Nevertheless, despite its popularity, database searching comes with some drawbacks^4^. The first problem with database searching is the computational complexity imposed when working with large databases. As the algorithm needs to consider all possible peptides derived from a protein sequence, the resulting databases will grow exponentially when taking into account multiple missed cleavages and a variety of potential post-translational modifications (PTMs)^3^. Another important disadvantage of database searching is the lack of peak intensity information and information on non-canonical fragments in the generated theoretical spectra, which limits the sensitivity of the scoring function.

Spectral library searching seeks to correct for these two issues, by comparing experimental spectra to a spectral library built from previously identified spectra^5^. Nowadays, this spectral library searching approach is gaining more attention due to a number of advantages^6^. Because the search space is confined to previously observed and identified peptides, the computational complexity is reduced^7^. Moreover, spectral searching can take advantage of all spectral features, including actual peak intensities and the presence of non-canonical fragment ions^8^, to determine the best possible peptide match. As a result, this technique often yields improved sensitivity^9^.

Different tools to apply spectral library searching have been developed over the past years, with SpectraST^10^, the National Institute of Standards and Technology (NIST) MS-Search^11^ tool and X!Hunter^12^ as notable examples. Each of these tools, however, comes with some limitations, such as a low processing speed, issues with handling large databases, and operational complexity. Furthermore, these tools typically support only specific spectral data formats, which also creates a threshold to adoption if the desired library is not presented in a compatible format. Taken together, these issues have prevented widespread adoption of the spectral library searching approach in proteomics.

We have therefore developed COSS (CompOmics Spectral Searching), a new, fast, and user-friendly spectral library search tool capable of processing large databases and supporting different file formats. Two scoring functions are available in COSS, namely MSROBIN, which relies on probabilistic scoring, and the cosine similarity score. COSS also offers an intuitive graphical user interface, allowing it to be adopted easily. To control the false discovery rate, a built-in mechanism to generate decoy spectral libraries has been provided as well. We have benchmarked COSS on three different spectral libraries and our results show that, compared to established tools, COSS delivers more reliable identification. At the same time, COSS requires drastically lower computation time and has a much-reduced memory footprint, eliminating the requirement for high performance and costly equipment, and further lowering the threshold to adoption of the spectral library searching approach.

## MATERIALS AND METHODS

### Implementation

COSS is developed in Java in a modular fashion so that its code is reusable and future-proof. Separate modules have been developed for key tasks such as indexing, filtering, matching, and decoy generation. To ensure maximal compatibility with input formats, the spectrum reader has been developed as a separate subsystem. COSS supports mzXML^13^, mzML^14^, ms2 and dta input formats through the mzIdentML^15^ library, while support for the msp and mgf formats is included through an in-house implementation. The compomics-utilities^16^ library was used for spectra visualization. Because COSS is completely developed in Java, it is platform independent, allowing users to run the software in their own preferred environment (e.g., Windows, Linux, or MacOS).

### Scoring function

COSS implements two scoring functions: MSROBIN, which is based on the probabilistic scoring function of Yilmaz et al.^17^, itself a derivative of the Andromeda scoring function^18^ and the cosine similarity score. The scoring procedure consists of two main steps. First, both the query and library spectrum are divided into 100 Da windows and within each window, the *q* peaks with the highest intensity are selected. Next, the score is calculated for *q* varying from 1 to 10 and the highest score is retained. The MSROBIN scoring function consists of two parts, an intensity part and a probability part. The probability scoring part is as follows:

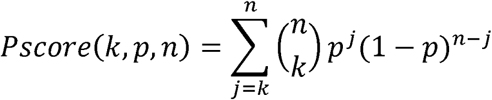

Where n is the number of peaks, *k* is number of matched peaks, and *p* is the probability of finding a match for a single matched peak, calculated by dividing the number of retained high intensity peaks by the mass window size which we set at 100 Da.

The second part is the intensity scoring which is calculated as:

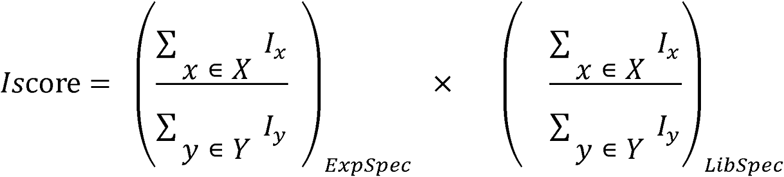

Here, *I* is the peak intensity, *X* is the set of matched peaks and *Y* is the set of intense peaks selected from each 100 Da window. The final score is then computed as:

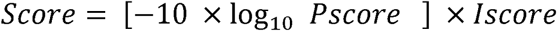

We have calculated the cosine similarity scoring function as follows:

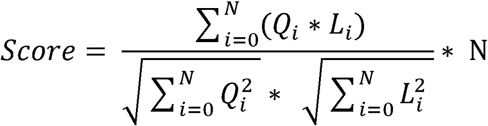

Where Q is the intensity of the matched peak found in the query spectrum, L the intensity of the matched peak found in the library spectrum and N is the total number of matched peaks between query and library spectra under comparison. The score is weighted by the number of matched peaks.

### False discovery rate estimation

Erroneous peptide assignments can occur due to poor spectrum quality or limitations in the scoring function. Validation of the obtained results is therefore a key step in peptide identification, and typically takes the form of false discovery rate (FDR) control^19^. For this purpose, COSS implements a decoy spectral library strategy, which can generate a number of decoy spectra equal to the size of the original spectral library using reverse and random sequence decoy generation technique as described in Zhang et al. ^20^. Briefly, the sequence of each spectrum is reversed, leaving the last amino acid in place. Based on this sequence, the masses of the *b, y* and *a* ions are calculated and the corresponding annotated peaks in the spectrum are moved on the m/z axis accordingly leaving the unannotated peaks in place.

The generated decoy spectra are concatenated to the original spectra in the library, and the search is run against this concatenated target-decoy spectral library. The corrected FDR value is then calculated as described previously in Sticker et al.^19^.

To evaluate whether the generated decoys accurately control FDR, we have used a modified entrapment method^21^. For this, we have added *Pyrococcus furiosus* spectra to the search data set, and then ran the search against the NIST library concatenated with the generated decoy spectra. FDR and false discovery proportion (FDP) are then calculated as follows:

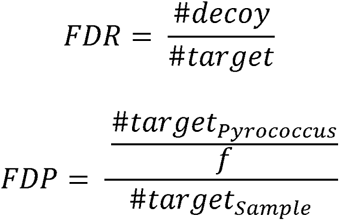

With *f* the fraction of *Pyrococcus furiosus* spectra over non-*Pyrococcus* spectra in the search.

### Benchmarking datasets and spectral libraries

We obtained raw data files from eleven runs from the Human Proteome Map^22^ (ProteomeXchange^23^ ID PXD000561) and ten runs from the deep proteome and transcriptome abundance atlas^24^ dataset (ProteomeXchange ID PXD010154) as benchmarking data sets (Table 1). All these 21 raw files were converted to Mascot Generic Format (mgf) format using the *msconvert* tool (ProteoWizard^25^), with the peak picking algorithm activated.

**Table 1.** Benchmarking data sets used to test COSS. The first eleven data sets are taken from the Human Proteome Map (ProteomeXchange ID PXD000561), while the last ten data sets are taken from the deep proteome and transcriptome abundance atlas (ProteomeXchange ID PXD010154). Individual raw files are selected randomly from their respective tissue.

| Raw file | Number of spectra | ProteomeXchange ID | Tissue |
| --- | --- | --- | --- |
| Adult_Adrenalgland_Gel_Elite_49_f22.raw | 14954 | PXD000561 | Adrenal Gland |
| Adult_Bcells_Gel_Elite_76_f01.raw | 7496 | PXD000561 | BCell |
| Adult_Colon_bRP_Elite_50_f24.raw | 1811 | PXD000561 | Colon |
| Adult_Gallbladder_Gel_Elite_52_f24.raw | 6146 | PXD000561 | Gallbladder |
| Adult_Heart_Gel_Elite_54_f05.raw | 12216 | PXD000561 | Heart |
| Adult_Kidney_Gel_Elite_55_f01.raw | 5267 | PXD000561 | Kidney |
| Adult_Liver_Gel_Elite_83_f23.raw | 4615 | PXD000561 | Liver |
| Adult_Ovary_Gel_Elite_58_f02.raw | 3885 | PXD000561 | Ovary |
| Adult_Pancreas_Gel_Elite_60_f02.raw | 2156 | PXD000561 | Pancreas |
| Adult_Retina_bRP_Elite_64_f15.raw | 1770 | PXD000561 | Retina |
| Adult_Spinalcord_Gel_Elite_67_f01.raw | 4105 | PXD000561 | Spinal cord |
| 01295_A01_P013198_S00_N01_R1.raw | 26621 | PXD010154 | Esophagus |
| 01349_A01_P013680_S00_N01_R1.raw | 31307 | PXD010154 | Placenta(ii) |
| 01087_C05_P010739_S00_N35_R2.raw | 38050 | PXD010154 | Stomach(i) |
| 01226_B05_P012502_S00_N34_R1.raw | 39053 | PXD010154 | Liver |
| 01276_G04_P013128_S00_N30_R1.raw | 43981 | PXD010154 | Stomach(ii) |
| 01347_E04_P013678_S00_N29_R2.raw | 47687 | PXD010154 | Pancreas |
| 01077_D03_P010695_S00_N20_R1.raw | 49460 | PXD010154 | Placenta(i) |
| 01698_G03_P018021_S00_N23_R1.raw | 50666 | PXD010154 | Pituitary/H. |
| 01349_D04_P013680_S00_N28_R1.raw | 50903 | PXD010154 | Placenta(iii) |
| 01075_D03_P010693_S00_N20_R1.raw | 51290 | PXD010154 | Colon |

Benchmarking was performed using three distinct spectral libraries (Table 2), obtained from NIST, PRIDE^24^ and MassIVE^25^.

**Table 2.** Spectral libraries used to benchmark COSS.

| Spectral Library | Total number of spectra | URL |
| --- | --- | --- |
| MassIVE<sup>25</sup> (Human HCD Spectral Library) | 2,154,269 | http://massive.ucsd.edu/ProteoSAFe/static/massive-kb-libraries.jsp |
| NIST (human HCD library) | 1,127,970 | https://chemdata.nist.gov/dokuwiki/doku.php?id=peptidew:lib:humanhcd20160503 |
| PRIDE Cluster<sup>24</sup> (Human) | 789,745 | https://www.ebi.ac.uk/pride/cluster/#/libraries |

### Running searches

All benchmarking is performed on the same virtual machine, equipped with dual Xeon E5-242016 processors at 1.90GHz, 28 GB of RAM, and running the Microsoft Windows 10 operating system. To run SpectraST, we used the Trans Proteomic Pipeline (TPP v5.2.0-b1) software for Windows. SpectraST was run in command line mode according to the user manual (http://tools.proteomecenter.org/wiki/index.php?title=Software:SpectraST). The spectral libraries, originally in msp format, were first converted to the splib file format, and then a consensus spectrum from the generated splib file was created. Quality control was applied on this consensus file, and finally decoy spectra were generated and appended to the consensus file. Search settings were a precursor m/z tolerance of 0.01 Th, with the rest of the settings left at their defaults (Supplementary methods).

Sequence database searches using MS-GF+^26^ (version v2018.04.09), have been performed through SearchGUI^27^ (version 3.3.16) and PeptideShaker^28^ (version 1.16.42). The search database is constructed from the human proteome (UP000005640) as obtained from UniProt^29^ (consulted on 9/10/2018). Carbamidomethylation of cysteine and oxidation of methionine are used as fixed, and as variable modification respectively. Trypsin is used as protease and a maximum of two missed cleavages was allowed. Precursor m/z tolerance was set to 10 ppm, and fragment tolerance to 0.05 Da. Precursor charges from 2 to 4 are considered.

To run COSS, decoy spectra were generated using the reversed sequence decoy generation technique. For the MassIVE and PRIDE libraries, where spectra are not (fully) annotated, we have first annotated the library using the built-in spectra annotator in COSS and a fragment tolerance of 0.05 Da. In this way, a, b and y ions where annotated, taking into account +1, +2 and +3 as possible charges and H2O and NH3 as possible neutral losses. The generated decoys were appended to the original spectral library and searches were performed using a precursor m/z tolerance of 10 ppm and a fragment m/z tolerance of 0.05 Da.

## RESULTS AND DISCUSSION

### Graphical user interface

COSS comes with a user-friendly interface that allows the user to set all parameters (Supplementary Figure S-1) needed for spectral similarity search. COSS supports most common MS/MS spectrum formats (including mgf, msp, ms2, mzML, mzXML, and dta). The user can generate decoy spectra for their spectral library using two types of decoy generation techniques that are implemented and integrated in COSS. COSS also provides an intuitive interface to visually inspect the obtained results (Figure 1). This interface reports all experimental spectra with matches in the spectral library in an interactive table, sorted by descending match score. When a query spectrum is selected, the top 10 matched spectra from the spectral library are displayed in the bottom table. For each match, the query spectrum and the matched library spectrum can be visually inspected. The results can be exported in tab-delimited text format, comma-delimited text format (CSV) and Microsoft Excel format (xlsx) for further processing and reporting. In addition to the graphical user interface, COSS also comes with a documented command-line interface to easily deploy the software on servers and high-performance clusters. The flexibility of COSS is further enhanced by its ability to run on all common operating systems.

**Figure 1.**
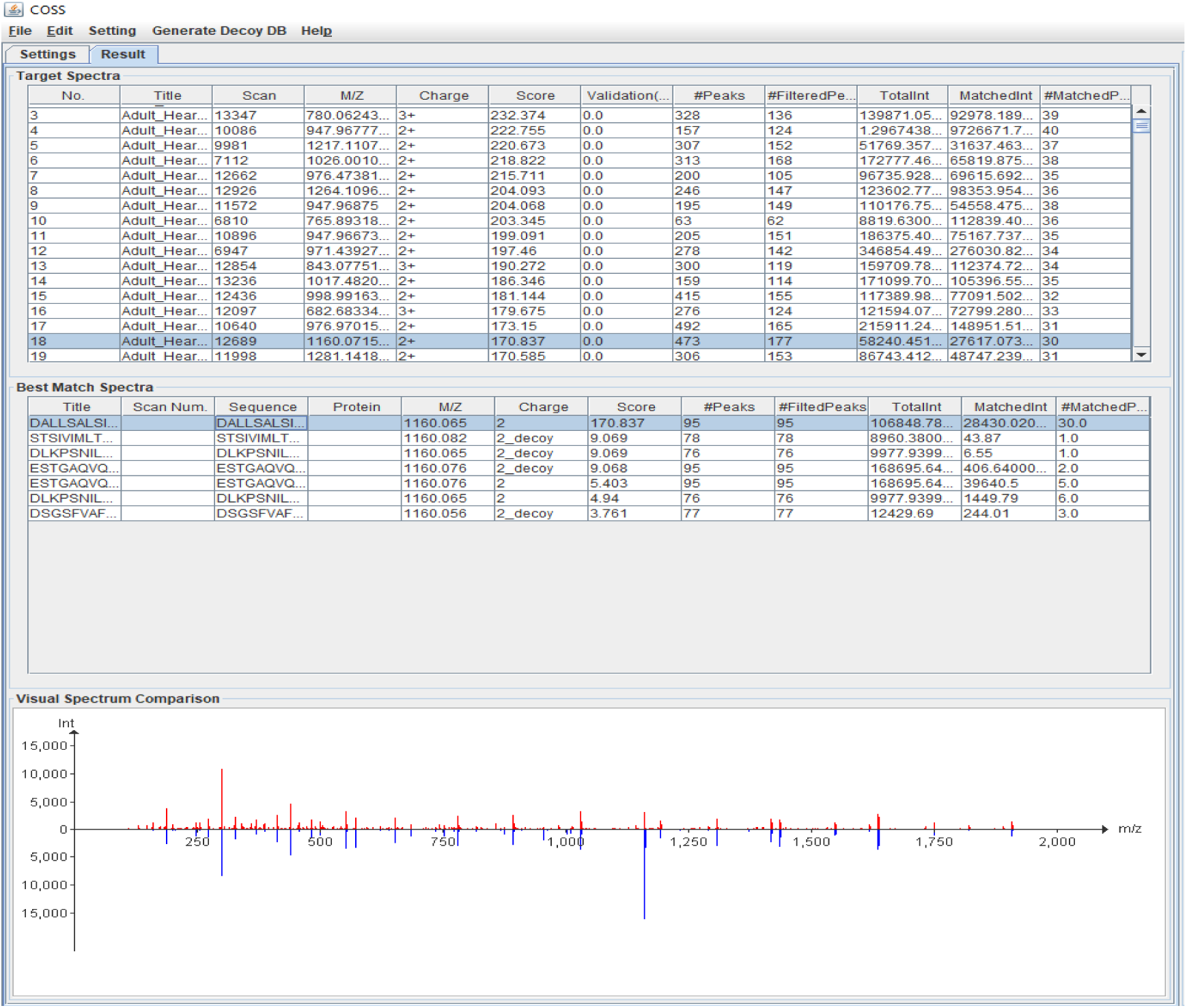
Search result interface of COSS: the upper table lists the experimental spectra while the lower table lists the top 10 matched spectra for the selected experimental spectrum. An interactive spectrum comparison view is presented at the bottom with the selected experimental spectrum (red) mirrored with the selected matched library spectrum (blue).

### Evaluation of false discovery rate estimation

To evaluate the ability of COSS to accurately assess the FDR based on the implemented decoy generation technique, we used the modified entrapment approach using *Pyrococcus furiosus* spectra^21^. Our results show that while SpectraST dramatically underestimates the FDR (at 1% estimated FDR, the actual FDP as measured by Pyrococcus identifications is 2.57%), COSS assesses it much more accurately (at 1% estimated FDR, actual FDP is 1.8%) (Figure 2).

**Figure 2.**
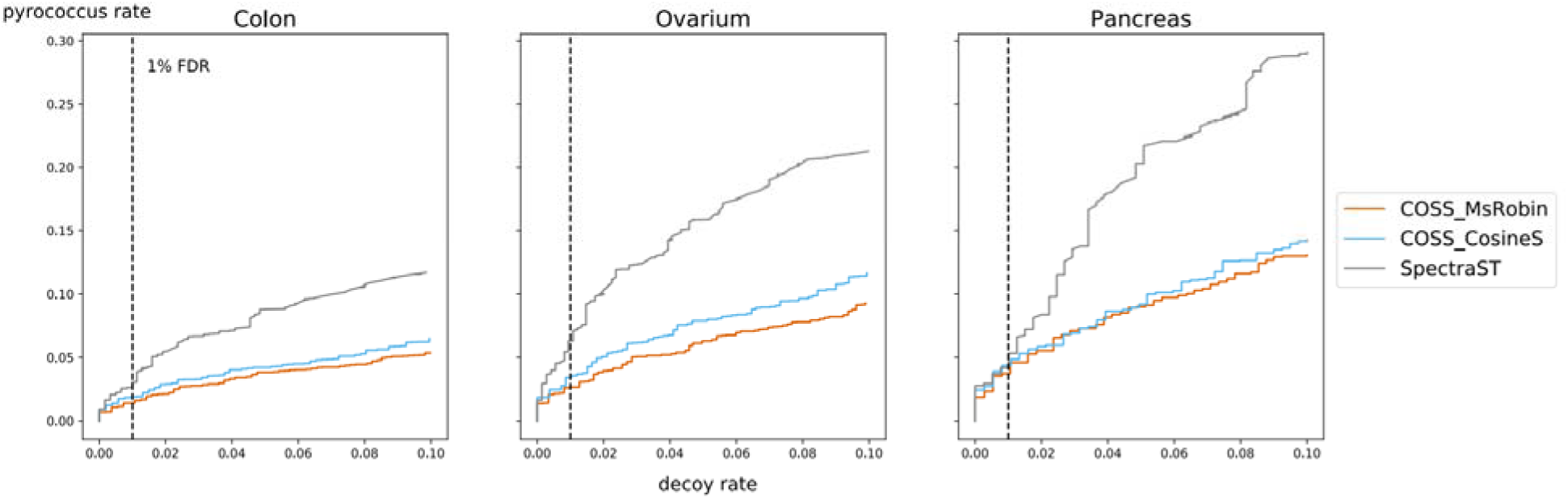
COSS more accurately controls the FDR. Here, decoy rate (FDR) is shown in conjunction with the *Pyrococcus* rate (FDP) for search results from three runs from the Human Proteome Map. Shown are the FDR and FDR from COSS with the MSROBIN scoring function, COSS with cosine similarity scoring function and SpectraST. The MSROBIN scoring function of COSS is the most reliable scoring function in terms of FDR estimation, narrowly beating the cosine similarity score, while SpectraST frequently underestimates the true FDR.

### Search result comparison

Figure 3 shows the identification rate of COSS (MSROBIN and Cosine similarity), SpectraST, and MSGF+. The overall identification rate of COSS on the first test dataset (PXD000561) are 13.4%, 9.4% and 16.1% against NIST, PRIDE, and MassIVE libraries, respectively, while the cosine scoring function of COSS obtains 12.7%, 8.8% and 15.5% against NIST, PRIDE and MassIVE, respectively. The identification results of SpectraST are 16.0% and 9.0% against NIST and PRIDE respectively. For the second test dataset (PXD010154), the identification rates are slightly lower, but the same trends are visible. Note that SpectraST’s performance against the MassIVE database could not be assessed as SpectraST could not handle a library of this size, even on our server with 28GB of RAM. MSGF+ identified 15.3% of the spectra when searching the human proteome database. As shown in Figure 3 the overall identification results of SpectraST, against NIST library, are slightly higher than those of COSS. However, considering our assessment of the false discovery rate estimation issues for SpectraST (Figure 2), the surplus of hits found by SpectraST are likely to be strongly enriched with false positives.

**Figure 3.**
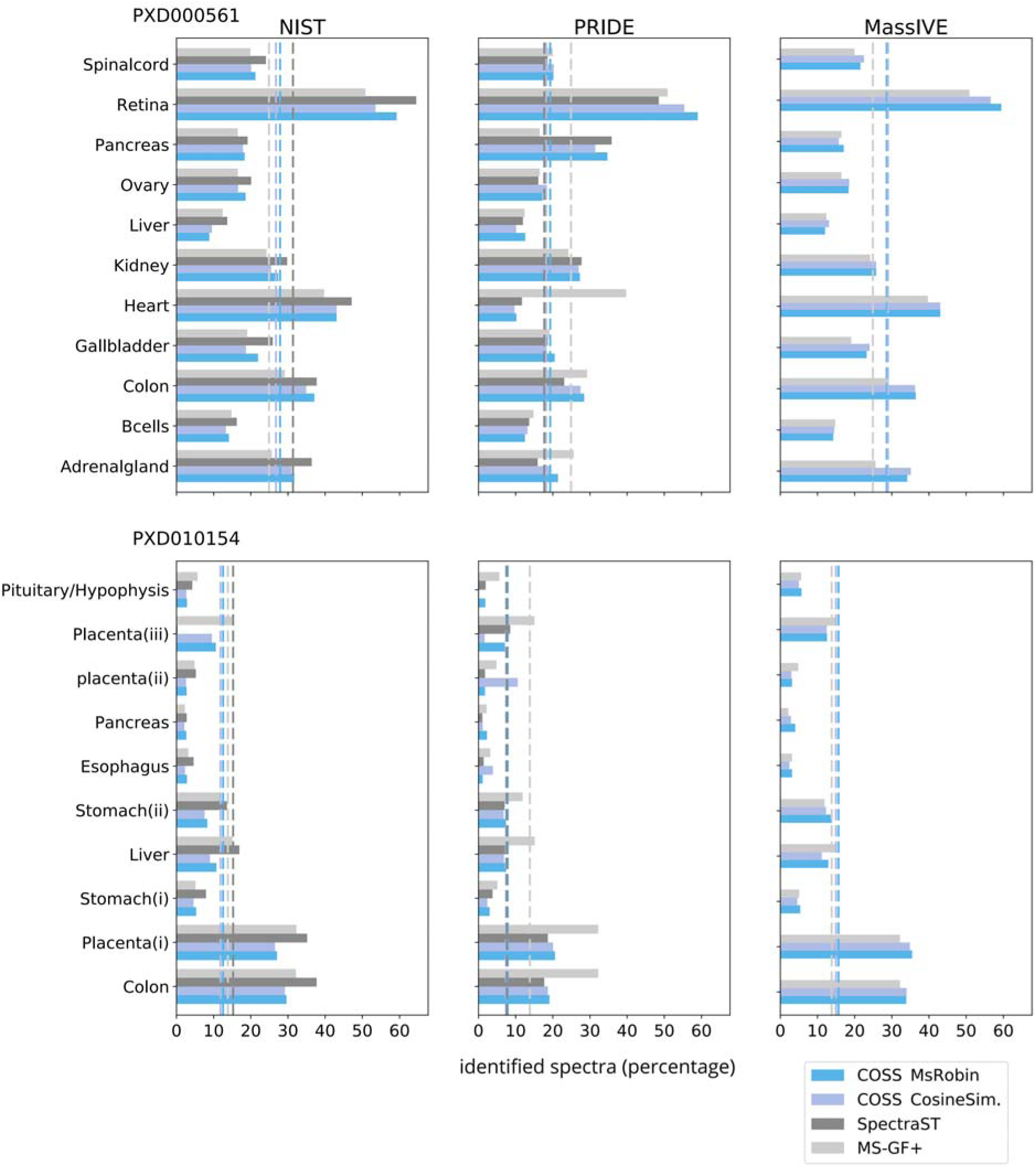
COSS performance evaluation against SpectraST and sequence database searching in terms of identification rate. Shown here is the identification rate against the NIST, PRIDE Cluster and MassIVE spectral libraries for COSS and SpectraST, and against the human proteome sequence database for MS-GF+. Due to excessive memory requirements, SpectraST could not run the MassIVE spectral library on our server with 28GB of RAM.

**Figure 4.xs.**
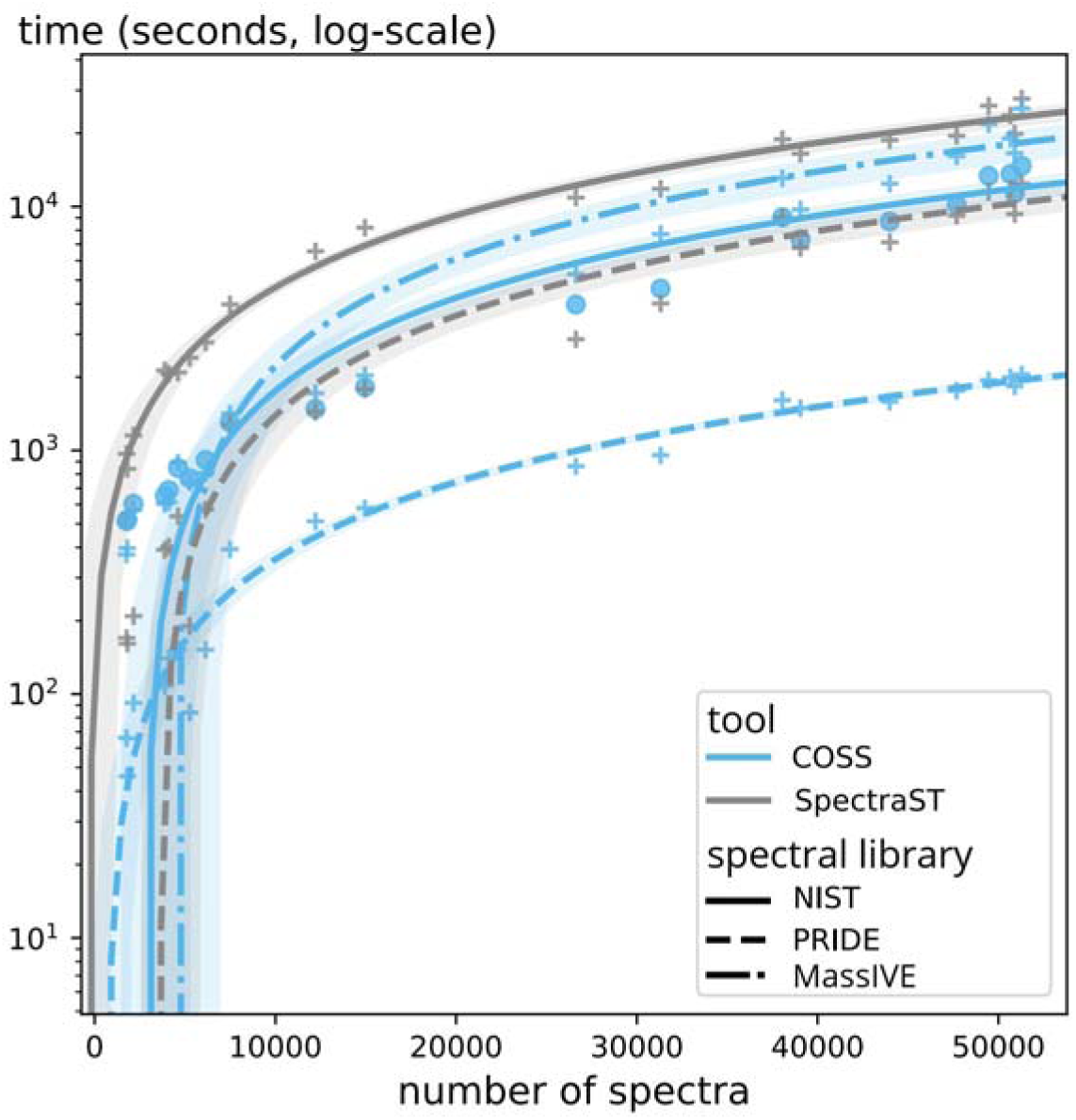
Execution times of COSS and SpectraST. While execution time increases with experimental data set and spectral library sizes, COSS clearly outperforms SpectraST in every case. Even for large data sets and spectral libraries, results for COSS are retrieved in well under half an hour.

The identification rate of COSS and SpectraST on the PRIDE Cluster spectral library are similar, however, lower than that of MS-GF+, most likely due to the incomplete coverage of this library (consisting of 189,400 unique peptides, compared to the 1,500,000 unique peptides derived from the *in silico* tryptic digest of the human proteome sequence database). In addition, the identifications of the three approaches show a good overlap in terms of the identified peptides (Supplementary Figure S-2). The identification overlap between the two algorithms of COSS is very large in all the three different libraries (Supplementary Figure S-3).

### Running time comparison

To evaluate the computational efficiency of the algorithm, we ran COSS and SpectraST on the same data sets using the same virtual machine and recorded the execution time for each algorithm. The results of the comparison are shown in Figure 3. While the size of the query dataset and the spectral library both clearly influence the executing time, we found that COSS drastically outperforms SpectraST in all cases. Here again, there is no data for SpectraST for the MassIVE library due to the inability to run SpectraST on this library on our server.

## CONCLUSIONS

There is a need for spectral library search tools that can easily analyse data from today’s high-throughput mass spectrometry-based proteomics experiments, and that can match tens of thousands of acquired spectra against proteome-wide spectral libraries. A few such search algorithms like SpectraST have already been developed but come with important limitations: long search times, an inability to handle very large spectral libraries, and limited input file format support. Here we present COSS, a user-friendly spectral library search tool that is fast, that can handle large datasets, and that supports the most commonly used MS/MS data formats. COSS offers both a graphical as well as a command-line interface, enabling users to perform anything from small-scale analyses on laptops to automated, large-scale data reprocessing on high-performance compute clusters. Because COSS is developed in Java, it is also platform independent, allowing it to run seamlessly on all commonly used operating systems. Furthermore, COSS’s modular architecture and open-source code invites and facilitates future development by the community at large. We have compared COSS to SpectraST and a sequence database search algorithm, MS-GF+, in terms of identification performance, and found that COSS offers a more reliable identification than SpectraST, while also drastically outperforming SpectraST in running time. While MS-GF+ can identify more spectra in a subset of data sets, the overall identification performance of COSS is quite similar. These properties make COSS highly suitable for large-scale analyses against ever expanding spectral libraries, including those that aim to cover an entire proteome of an organism.

## AVAILABILITY

The software and its source code can be freely downloaded from https://github.com/compomics/COSS and is licensed under the permissive, open-source Apache License, version 2.0.

## ACKNOWLEDGMENTS

This project is supported by the National Institute of Health (NIH) [NCI-ITCR grant number 1U24CA199347 to G.A.S.], Research Foundation - Flanders (FWO) [3E023815 to E.V., grant number 1S50918N to R.G., grant number G042518N to L.M.], the Horizon 2020 programme of European Union project EPIC-XS [grant number 823839], and Kom op tegen Kanker (Stand up to Cancer), the Flemish cancer society [to P.V.]. We would like to thank Zheng Zhang from NIST, USA for providing us with the R code for generating decoy spectra libraries. We would also like to thank all the CompOmics group members for their ideas, discussions and support.

