## Supplemental file for "COSS: A fast and user-friendly tool for spectral library searching"

SUPPLEMENTARY FIGURES

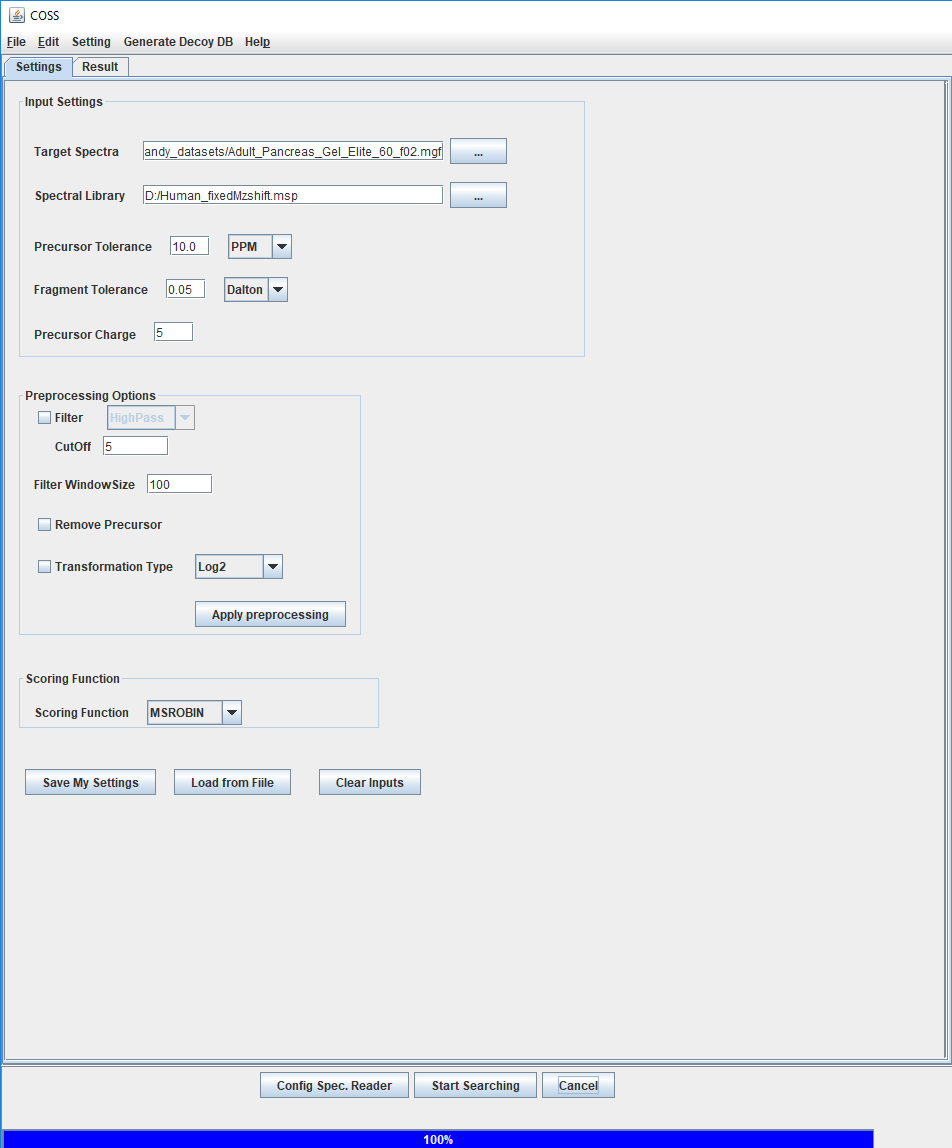

**Figure S-1.** COSS’s graphical user interface. Settings tab lets users to set search parameters and input files. The lower panel contains buttons to configure the spectrum reader and to start the and stop the search. It is also possible to generate decoy spectra for the given library by selecting a decoy generation method from the “Generate Decoy DB” menu.

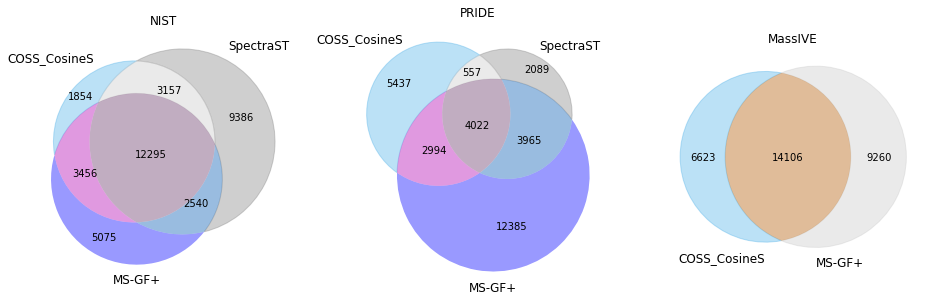

**Figure S-2.** Peptide level result overlap of the identifications from COSS, SpectraST and MS-GF+ against the NIST, PRIDE and MassIVE spectral libraries. The overlap between the search tools is more in the case of NIST library while the overlap is very less in the case of PRIDE would most likly be due to the small size of this library.

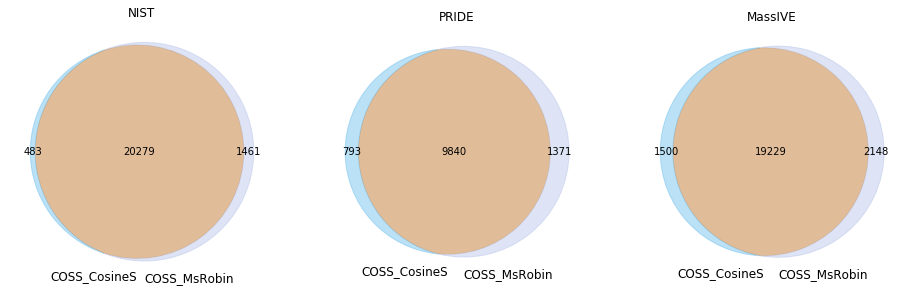

**Figure S-3.** Identification overlap between the two algorithms of COSS. Both MsRobin and cosine similarity algorithms of COSS overlap nicely in all the three libraries.

Supplementary methods

SpectraST configuration file:

```
### COMMON OPTIONS ##############################

# Spectral library to be searched against - required if not specified on command-line with -sL option - replace with your own
libraryFile = C:/TPPdata/cons_human_hcd_selected_QC_Decoy.splib

# A database file to be included in .pepXML outputs for downstream TPP processing - replace with your own
#databaseFile =

# The type of the database: "AA" = amino acid database; "DNA" = nucleotide database
databaseType = AA

# Whether to keep all accessed library entries in memory
indexCacheAll = false

# Precursor m/z tolerance
precursorMzTolerance = 0.01

# Output file format: xls (tab-delimited text)
outputExtension = xls

# Output directory: where to put the output files. Defaults to the same directory as the search data file.
outputDirectory = C:/TPPdata/SpectraST_SearchResult
```
